# Deep Learning of Cellular Metabolic Flux Distributions Predicts Lifespan

**DOI:** 10.1101/2024.11.22.623650

**Authors:** Tyler A.U. Hilsabeck, Shane L. Rea

## Abstract

It is a common observation that individuals within a species age at different rates. Variation in both genetics and environmental interaction are generally thought responsible. Surprisingly, even genetically identical organisms cultured under environmentally homogeneous conditions age at different rates, implying a more fundamental cause of aging. Here we have examined the basis for lifespan variance in haploid, single-celled yeast of *Saccharomyces cerevisiae*. The probabilistic nature of metabolism means metabolites often, but not always, follow the same route through the metabolic network. We speculate redundancy in metabolic pathway choice is sufficient to explain lifespan variance. To interrogate the reaction flux space of *S. cerevisiae* we used a model of its intermediary metabolism, comprising 1,150 genes, 4,058 reactions, and 2,742 metabolites (yeast GEM_v8.5.0). We restricted traffic through the metabolic network by knocking out each of the 1,150 genes, then generated a total of 406,500 flux distributions spanning the solution space of the resulting 812 viable mutants. We used replicative life span (RLS) data for the 812 viable mutants, corresponding to 66,400 individual cells. Four approaches were then employed to test whether reaction flux configuration could be used to predict lifespan: Principal Component Analysis (PCA) in conjunction with non-linear modeling of RLS; deep learning of RLS using either a Regression Neural Network (RNN) or a Classification Neural Network (CfNN); and deep learning using a convolutional neural network (CNN) following conversion of flux distributions to pixelated images. The four approaches reveal a core network of highly correlated reactions controlling aging rate that is sufficient to explain all lifespan variance. It includes biosynthetic pathways encompassing ceramides, monolysocardiolipins, phosphoinositides, porphyrin and glycerolipids. Our data lead to two novel conclusions. First, variance in the replicative lifespan of *S. cerevisiae* is an emergent property of its metabolic network. Second, there is convergence among metabolic configurations toward three meta-stable flux states – one associated with extended life, another with shortened life, and a third with wild type life span.

**One Sentence Summary:** Traffic routes and rates through the metabolic network of *S. cerevisiae* fully account for variance in replicative lifespan.

## INTRODUCTION

Age is a primary risk factor for debilitating diseases such as Alzheimer’s Disease, Parkinson’s Disease, cancer, cardiovascular dysfunction, and osteoporosis1–5–. In recent years, significant effort has identified conserved genetic- and biochemical processes that modulate the rate of aging in a variety of organisms. Despite such progress, our understanding of aging and how it impacts lifespan and disease incidence remains far from complete. In broad strokes we know that aging occurs at different levels in an organism: At a subcellular level, we know, for example, that key quality control processes such as protein turnover and DNA repair become increasingly compromised with time. At a cellular level, terminally differentiated cells such as neurons are lost without replacement. And, at a systemic level, we know that some tissues are likely to fail before others - the gut and vasculature, for example, are age-dependent weak points in the environmental barrier of humans. As one organ begins to fail, stereotypical compensatory responses are activated, but one damning sequelae of aging is that compensatory robustness narrows with time (homeostenosis).

Less well appreciated in the process of aging is the role of stochasticity. This is best illustrated by considering a population of genetically identical individuals cultured under environmentally homogeneous conditions – not all individuals die at the same time. Indeed, survival curves for such organisms are usually just scaled versions of those measured for higher organisms. Stochasticity comes in many flavors: At a quantum level, thermal fluctuations place atoms and molecules in different energetic states. In cells, thermal motion places molecules at random points in space. These probabilistic events are almost impossible for us to control but their effects are felt whenever life depends on a countable number of molecules, such as, for example, fate choice by transcription factors. Higher up the stochastic ladder, we know that unwanted chemical modifications accumulate with time, and these events almost certainly underlie some of the reductions in the overall efficiency of protein turnover and DNA repair that was alluded to earlier. Higher up still, we know that there are certain kinds of chance events that potentially greatly affect organismal aging, but which can be controlled to some extent. Flux through the reactions of intermediary metabolism is, in every sense, a probabilistic process and metabolites are confronted with great choice inside cells. Cells can lower this choice by placing weights on different reaction pathways – that is, by controlling the expression level of catalysts, by controlling the subcellular localization and co-assembly of such catalysts, and through metabolite shuttling where compounds are force-fed between related catalysts. In this way, stochasticity can be reduced.

In this study we examined the extent to which choice of flux route through the entire network of intermediary metabolism is a significant predictor of lifespan outcome in clonal populations of the single-celled yeast, *S. cerevisiae*. Lifespan in this species can be followed by measuring the capacity of individual mother cells to generate progeny via asymmetric budding (termed replicative lifespan^7^). We find that redundancy in how compounds are fluxed through metabolic pathways is sufficient to account for all variance in *S. cerevisiae* lifespan.

## RESULTS

### Defining the Metabolic Solution Space of S. cerevisiae

The metabolism of *S. cerevisiae* can be represented by a multi-dimensional space (*n-space*), with flux through each reaction denoted as a dimension. Points in this space correspond to metabolic configurations, and those specific points that lead to no net change in intracellular metabolite concentrations while metabolites flux through the metabolic network correspond to positions of steady-state equilibrium. Due to physicochemical and biological restraints, the flux potential of the metabolic network is a bounded space. To determine whether there is a relationship between position in metabolic *n-space* and lifespan (**Fig. 1a**), we utilized the yeast consensus genome-scale model (GEM)^8^, version 8.5.0^9^, which combines 4,058 biochemical reactions across 14 sub-cellular compartments, 2,742 metabolites, and 1,150 genes, to provide an *in silico* description of *S. cerevisiae* (*strain S288c*) metabolism^8^. The 1,150 genes encode enzyme catalysts and metabolite transporters and represent ∼1/5 of the yeast genome (https://www.uniprot.org). Knockout mutants of metabolic genes naturally limit the metabolic configurational space of wild type yeast. Such mutants provide a tool by which different regions of wild type configurational space can be probed to determine their relationship with lifespan. To interrogate how removing genes from *S. cerevisiae* restricts its network configurational space, we used constraint-based modeling^10^. We limited our analysis of Yeast GEM v8.5.0 to aerobic growth on glucose under defined minimal media conditions and used flux balance analysis (FBA) to identify network-wide reaction fluxes that placed the metabolic network into a steady state equilibrium when maximizing biomass production, a proxy for yeast growth. Using this approach we identified 812 viable yeast knock-out mutants that had non-zero biomass production values. Since the stoichiometric matrix for the metabolic network in *S. cerevisiae* is underdetermined, flux variability analysis (FVA) was used to chart the boundaries of the FBA solution space for each viable mutant (**Fig. 1b)**. The resulting FVA-constrained flux space (**Fig. 1c**), although bounded, effectively contains an infinite number of solutions. We therefore used Monte Carlo sampling (**Fig. 1d**) to identify a set of configurations for each knockout mutant that evenly spanned their flux space. We then removed reactions across all configurations that had either zero variance across all mutants or had variances less than 10^−10^ flux units (**Fig. 1e, f**). This resulted in a total of 406,500 flux configurations among a reduced set of 2,912 reactions that spanned the combined solution space of all 812 viable single-gene mutants.

### Replicative Life Span

Replicative lifespan (RLS) employed was used undertaken manually using microdissection for all 812 viable knockout mutants{Formatting Citation}. A total of 66,393 individual cells were followed until replicative exhaustion, with approximately equal numbers of mutant and pair-matched wild type cells measured during every assay. The number of individual cells for which RLS measurements were recorded varied by knockout and ranged from five to 1,717 (**Fig. 1g**). The comprehensive wildtype RLS dataset (n=36,000) established the expected range of yeast lifespan (**Fig. 1h**). Mutants with averages outside these parameters showed biologically meaningful divergence (p <0.0035) from normal cellular aging trajectories. If metabolic flux distribution is predictive of RLS, then we anticipate the RLS of any knockout mutant should be no larger or no smaller than the most extreme wild type RLS values, since the metabolism of each mutant is a subset of wild type. Indeed, RLS values of all knockout yeast fell within the span of wild type yeast RLS values (**Fig 1i**, normalized RLS values (ΔRLS) are shown). Since we were unable to assign ΔRLS values to individual configurations within the flux space of each mutant, we associated each mutant’s average ΔRLS value to all its configurations. Mutual information calculations showed that all 2,912 reactions with variance across the 406,500 flux distributions held lifespan-related information, but this information content was not distributed evenly (**Fig. 1j**). In subsequent sections we extracted this information to predict yeast survival. A summary of the metabolic and RLS data employed for this study is provided in **Fig. 1k**.

### Principal Component Analysis and Non-Linear Modeling of ΔRLS

To determine if position in metabolic configuration space can predict ΔRLS, we first used Principal Component Analysis (PCA) in conjunction with a Generalized Additive Model (GAM). PCA was used to further reduce the dimensionality of configurational space - from 2,912 reactions to 526 Principal Components (PCs), where each PC is comprised of a weighted contribution (loading score) of each of the original reactions. 99% of the data variance was captured by these 526 PCs (**Fig. 2a**). We next utilized a Generalized Additive Model to fit ΔRLS values using all 526 PCs, but this approach failed for three reasons – first, we had insufficient computational power to test the full model (812 ΔRLS values, 406,500 data points and 526 PCs); second, the frequency distribution of ΔRLS values was not even (**Fig. 2b**), meaning the GAM focused heavily on fitting the most frequent data; and finally, assignment of a mean ΔRLS value to all 500 configurations of each mutant confounded the spline fitting procedure because configurations that overlapped in n-space were assigned more than one ΔRLS value. We therefore refined our PCA/GAM modeling approach, first by testing whether reactions with only the highest flux rates (high flux backbone, refer to **Fig. 1e**) held ΔRLS-differentiating information (they did not), and then by modeling only the centroid configuration of each mutant. Also, we binned centroids to weight the data more evenly before GAM fitting (**Fig. 2c**). Finally, to overcome the computational limitation, we established an algorithm (*GAMRandomStep)* that allowed us to both identify PCs with predictive power (while only screening a fraction of the PCs at any one time) and to find the optimal number of PCs that maximized the fitting of each model (**Fig. 2d**). In the absence of binning, the best centroid-fitting model identified by *GAMRandomStep* was comprised of 8 PCs and used a search size of twenty-five PCs per iteration (hereafter referred to as PCA_cen). 10,000 randomizations of the ΔRLS values among the rotated centroid configurations showed the amount of deviance explained by this model (18.8%) was significant at 9.34σ (**Fig. 2e**). Although this model contained ΔRLS-predicting information (**Fig. 2f**) it performed less favorably when attempting to predict ΔRLS for all 406,500 configurations (**Fig. 2g**). Binning of the centroids led to increasingly better fitting models (**Fig. 2c**); with 100 evenly spaced bins (resulting in 71 occupied bins, **Fig. 2h**) yielding the two best fitting models (**Fig. 2c, i, j**). The amount of deviance explained by the best fitting GAM was 83.8% (using 6 PCs). This model’s predictive value partially transferred when tested on all 812 centroids (**Fig. 2k**). The splines generated by the GAM for this model, and their partial predictions of ΔRLS, are shown in **Fig. 2l** and **Fig. 2m**. Finally, the loading scores for each reaction in the *second-best* fitting GAM (71 bins, 5 PCs, 79.0% explained deviance) are presented in **Fig. 2n**. We have presented data for this model (hereafter referred to as PCA_bin), and not the best fitting model, to illustrate a key point: both models utilize different, albeit overlapping, sets of PCs, even though they have the same approximate predictive power. Approximately 20% (600) of the reactions on each PC have non-zero loading scores and so presumably the most important reactions controlling ΔRLS are still equally covered by both sets of PCs. To further refine the reactions that control ΔRLS we noted that PC loading scores often fell into three classes, distinguishable by absolute magnitude (**Fig. 2n**, ***top row)***. We expected those reactions with the biggest difference in flux when comparing the averages of knockouts representing the top and bottom 5% of ΔRLS values to also have the greatest absolute loading score values, but this was not the case (**Fig. 2n**, ***middle row***). We therefore weighted the magnitudes of the loading score of each reaction in each PC by the average difference in reaction flux rate between mutants with the top versus bottom 5% of ΔRLS values, and subsequently applied a heuristically-determined cutoff to extract a set of 593 unique reactions (**Fig. 2n**, ***bottom row***). We presume that some or all these reactions drive changes in ΔRLS.

### Modeling of ΔRLS Using Neural Networks

Neural networks provide an alternate approach for non-linear data modeling. An advantage of these algorithms is that input variables remain independent, meaning their importance to the accuracy of the final trained model can be determined by simply excluding them. Three classes of neural networks were employed to identify reactions controlling the rate of aging in *S. cerevisiae.* First, we attained limited success using regression neural networks (RNN). Specifically, after testing several thousand models in which we systematically varied the network architecture (neuron size and connectivity) and multiple hyperparameters (activation function; regularization method including neuron dropout and k-fold cross validation; batch size; learning rate; number of epochs; number of input neurons (reactions); and a custom weighting parameter that forced the modeling to focus on specific subsets of data), we concluded that an RNN could not accurately model all 406,000 mutant flux configurations. It is possible that the same limitations that negatively impacted the PCA/GAM modeling were responsible for poor RNN modeling performance, namely the frequency distribution of the ΔRLS values and our assignment of a single ΔRLS value to all 500 configurations for each mutant. However, we succeeded in using an RNN to accurately predict ΔRLS using just the centroid configurations of all 812 knockouts. Two models were particularly effective - the first employed a stacking approach where three separate RNN models were generated to fit centroid flux configurations across neighboring ΔRLS ranges. This model employed the 526 reactions with the highest-ranking mutual information scores as input variables and fit the data with an RMSE of 0.574. This model could also successfully predict individual as well as binned centroid values (**Fig. 3a**). The second RNN model used the top 536 reactions with the highest-ranking mutual information scores, 10-fold cross-validation, but no stacking. This model had average R^2^ and RMSE values of 0.99998 and 9.43E-07, respectively (**Fig. 3b**, *green box*). By individually excluding each reaction from this RNN, we identified 160 reactions responsible for segregating the centroid flux configurations among the 812 knockout mutants (**Fig. 3b**, *green box, bottom panel*).

### Modeling of ΔRLS Using Classification Neuronal Networks

Our inability to build an RNN that could reliably distinguish all 406,000 configurations of the 812 viable knockout mutants might reflect the limitations we just mentioned, or perhaps another factor is at play. Up to this point, we have hypothesized that every metabolic configuration is associated with its own ΔRLS value (even though in practice we were forced to assign one ΔRLS value to all 500 configurations of each mutant). This hypothesis ignores the possibility that metabolic configurational switching in cells might only occur following large flux excursions in one or more reactions. So instead, it could be that a group of configurations ‘resonate’ around a central flux configuration (that represents a local minimum of chemical potential energy, say), and that configurations belonging to each group all lead to the same ΔRLS. Under this scenario, configuration switching between groups would require substantive energy input, and groups of configurations and their associated ΔRLS values would appear quantized. This idea is analogous to multi-stable states in chemistry (**Fig 3c**)^11^.

Replicative lifespan curves, like the survival curves they emulate, are inverted sigmoid in shape because the distribution of individuals that gives rise to them is bell-shaped (**Fig. 3d**). The steep walls of the bell curve could be interpreted as most individuals in a population ‘trying to be the same’. In other words, for *S. cerevisiae*, it is possible there is one major metabolic configuration that every cell in the population attempts to maintain when confronted with the same environment. The ends of the bell distribution might therefore represent individuals which have inadvertently found themselves in tangible new configurations. If it is true that groups of metabolic configurations behave as if quantized for ΔRLS, then modeling ΔRLS using them becomes a classification problem rather than one of regression.

Since we have no idea of the number of configurations that could potentially stably exist on the *S. cerevisiae* metabolic network, or indeed of the reaction flux boundaries that separate them, we explored this possibility by simply grouping the collection of 406,000 knockout configurations into ‘positive’, ‘zero’ and ‘negative’ ΔRLS clusters. We utilized a custom ΔRLS threshold grouping parameter to adjust and scan the boundaries between the three clusters, and in this way, we systematically tested several thousand classification neural networks (CfNNs) for their ability to accurately distinguish yeast knockouts and predict their replicative lifespan potential. We identified three notable sets of 5-fold cross-validated CfNN models (**Figs. 3e-g**). The first set (Set A) was trained on 197,056 flux configurations drawn predominantly from knockouts with the highest and lowest 30% ΔRLS values, and it used 526 reactions with the greatest mutual information scores as modeling variables. Set A models had an average accuracy of 87.4% (+/-0.03 SD) when tested on 49,264 unseen configurations (**Fig. 3e**). The second group of models (Set B) also used 526 reactions with the greatest mutual information scores as modeling variables, but was trained on 35,380 configurations drawn predominantly from knockouts with the highest and lowest 5% ΔRLS values. Set B models had an average test accuracy of 93.8 % (+/-0.005, SD) when tested on 8,845 unseen configurations (**Fig. 3f**). Finally, the third set of models (Set C) was like Set B, except all 2009 reactions with non-zero mutual information scores were used as modeling variables, along with some other minor differences (**Supp Table 2**, summary file). Set C models had an average test accuracy of 85.4 % (+/-0.194, SD) when tested on 8,845 unseen configurations. Shown in the middle and bottom rows of **Figs. 3e-g** are the k-fold model with the highest test accuracy. Quite remarkably, the search routine that we employed to optimize our custom ΔRLS threshold grouping parameter, and which resulted in these three, best-performing sets of CfNN models, independently converged in all three instances to define the boundaries for the ‘positive’ and ‘negative’ ΔRLS clusters as the flux configurations above the 95^th^ percentile and below the 5^th^ percentile, respectively (**Figs. 3e-g**, *top row*).

We increased the accuracy of our CfNN modeling predictions further by combining the individual predictions of the five models in Set B into a ‘majority vote wins’ ensemble (hereafter referred to as the ‘526_CfNN’ ensemble). We did the same also for Set C (hereafter called the ‘2009_CfNN’ ensemble). Using this approach, prediction accuracies increased to 95.0% and 97.0%, respectively. To identify reactions that were driving each ensemble’s accuracy, we individually removed the 526 or 2009 reactions used as modeling variables, and then re-determined each ensemble’s prediction accuracy. For the 526 CfNN ensemble there was a total of 115 reactions that negatively impacted accuracy, while for the 2009 CfNN ensemble there were 192 reactions (**Fig. 3h**). 78 reactions were common (**Fig. 3i**). No reaction, when removed, had an impact size on ensemble accuracy that exceeded 8%.

### Modeling of ΔRLS Using Convolutional Neuronal Networks

Encouraged by the ability of CfNN classifiers to predict ΔRLS for thousands of metabolic configurations, we tested the performance of Convolutional Neural Networks (CNNs) on the same task. CNNs excel at classifier tasks requiring image recognition. We therefore converted all 43,500 flux configurations from knockouts with the highest and lowest 5% ΔRLS values into images. This was accomplished using the Image Generator for Tabulated Data (IGTD) algorithm^12^. Flux values were range scaled across the 43,500 flux configurations to 256 grey scale using double-precision, floating point numbers. The IGTD algorithm assigns reactions to pixel positions in a matrix of chosen size so that the resulting correlation coefficients between all reactions (in this case across the 43,500 flux configurations) best emulates the physical distance between all pairwise pixel combinations (**Fig. 4a**). We also established a novel metric, called the ‘Inter-InterQuartile Range (IQR)-to-Median Difference Ratio’ (IIMDR, **Fig. 4b**) that facilitated assessment of the quality of our neural net modeling. IIMDR values range from zero to one, where positive values indicate two medians being compared differ at the 5% significance level. The IIMDR metric incorporates both variance and median difference measures, such that a score of zero indicates the two medians do not differ significantly, while scores close to one indicate non-overlapping medians and minimal variance in both data sets. In this way the *certainty* of the difference between two medians is emphasized at the expense of de-emphasizing the *magnitude* of the difference. We calculated the median flux of every reaction in the 2912 reduced reaction set across all 22,500 configurations of the 45 knockouts representing the bottom 5^th^ ΔRLS percentile, and likewise across all 21,000 configurations of the 42 knockouts representing the top 5^th^ ΔRLS percentile. Using these values, IIMDR scores were prepared for each reaction (**Fig. 4c**).

We assessed a total of 1,260 CNN models in which we systematically altered four hyperparameters (batch size, number of filters, kernel size and regularization technique). Parameter screening differed from our earlier approaches. Instead of using k-fold cross-validation, this time 14 independent searches were undertaken, all using the same set of hyperparameter choices, but differing by the seed used to initialize the random number generator, which in turn was used to initiate the neuron weights at the beginning of the optimization routine. All models were trained on 29,000 flux configurations and tested on 14,500 unseen flux configurations. All hyperparameter choices, except two, resulted in *median* test accuracies no better than guessing (**Fig. 4d**). The best choice of hyperparameters resulted in 14 models with a median test accuracy of 76.4%. Within this group, the top five models had a *mean* test accuracy of 87.2% (+/− 0.07, SD). Notably, for almost every set of hyperparameter choices tested, at least one CNN model performed far above its relevant median reflecting the importance of the initial neuronal weights in ultimately finding useful models.

To increase the accuracy of our CNN modeling we once again adopted a ‘majority vote wins’ approach and established two ensembles comprising multiple models. The first was composed of five models, all established using the same set of hyperparameters (CNN_A), and the second of seven models representing the best-performing models across all 1,260 CNN models constructed and each having a test accuracy >99% (CNN_B, **Fig. 4d**, *blue* and *red arrows*, respectively). Using this approach, test accuracies increased to 96.69% and 100%, respectively. Treating each ensemble separately, and by individually removing pixels (reactions) from the starting test images, we identified reactions controlling the ability of each ensemble to correctly predict ΔRLS values using flux configurations belonging to knockouts from the top and bottom 5^th^ percentiles. Focusing on the most powerful ensemble first, CNN_B, 395 reactions decreased prediction accuracy when individually removed (effect size > 0%). All 395 reactions are presented in log-transformed form in **Fig. 4e** (see also **Fig. 4i**, *top row*). Notably, 128 of these reactions exceeded 5% effect size, with the biggest reduction in accuracy reaching 45%. These reactions are presented without transformation in **Fig. 4f** (see also **Fig. 4i**, *second row)*. Evidently, not only are the 128 reactions differential for the two sets of knockout mutants, their position in the upper right corner of the reaction location map corresponds to the red arrows shown in the reaction correlation matrix of **Fig. 4a** and implies this group of reactions are strongly anti-correlated with another part of metabolism. We identified an almost identical set of reactions driving the prediction success of the second ensemble, CNN_A.

There are 230 reactions within the Yeast_v8.5.0 GEM that are directly impacted when all the genes encoded by the 87 knockout mutants belonging to the top and bottom 5 ^th^ percentiles of ΔRLS values are removed. Given the complexity associated with neural networks^13^, it is possible these 230 reactions were simply assigned their own pixel during CNN modeling, and this accounts for the striking accuracy of our ensembles. Considering first the entire set of 395 reactions that have non-zero effect sizes on CNN_B accuracy (**Fig. 4e)**, 42 of the 230 reactions impacted by the 87 KOs indeed overlap (**Fig. 4g**). This number is more than the 35 expected by chance (*hypergeometric probability, 0.008*), but far fewer than 230. Considering next only the 128 reactions with effect sizes > 5% (**Fig. 4f)**, the number of overlapping reactions is 10. This number is not different from chance expectation alone (*hypergeometric probability, 0.133*). Of these 10 reactions only one corresponds to a knockout gene that exclusively targets a single reaction which, given there are 19 such genes among the 87 total (**Fig. 4h**), this number is also not different from chance expectation (*hypergeometric probability, 0.373*).

### Convergence of RNN, CfNN and CNN Models

We have identified seven models, or ensembles of models, that vary in their relative success for predicting yeast ΔRLS based on metabolic flux configuration. PCA_cen and PCA_bin used centroid configurations and a small number of principal components to predict ΔRLS. These models resulted in an explained deviance of 18.8% and 79%, respectively. The RNN_536 model used centroid configurations in conjunction with 160 discrete reactions to predict ΔRLS, and this model fit the data with an RSME of 9.43E-07. For the CfNN_526, CfNN_2009, CNN_A and CNN_B ensembles, we employed tens of thousands of individual flux configurations to model ΔRLS, either from yeast with ΔRLS values in the top and bottom 30^th^ percentiles (CfNN), or just from the top and bottom 5^th^ percentiles (CNN). The resulting prediction accuracies of these four models on validation data were 95%, 97%, 96.7% and 100%, respectively. Beyond the obvious differences in success rate and data subsets used for the seven models, there are two other striking differences among the seven models in relation to the predictions they make. First, as we have already seen, the reactions driving the accuracy of the ΔRLS predictions are not always the same (compare **Fig. 3h** with **Fig. 4f**). Second, the relationship between flux configuration and ΔRLS that the models predict are of two types. Explicitly, PCA_cen, PCA_bin and RNN_536 all predict a continuous function, while the four ensembles predict a quantized function where flux configurations effectively condense to just three values: positive, zero (wild type) and negative ΔRLS. How might these apparently contradictory findings be reconciled?

The left most panel of **Fig. 5a** shows all 160 reactions responsible, to varying degrees, for accuracy of the RNN_536 model. The reactions have been plotted onto the post-IGTD reaction location matrix, which was also used to display the important reactions of the CfNN- and CNN ensembles shown in **Figs. 3h** & **4f**, respectively. Shown also in **Fig. 5a** are the Olden Importance scores for each of the 160 plotted reactions. There is a cluster of 26 reactions with strongly positive Olden Importance scores (green), meaning they largely drive success of the model. To determine if the RNN algorithm naturally grouped knockouts in its bid to locate a model that could accurately predict ΔRLS, we calculated IIMDR values for all 160 reactions using binned centroid fluxes from increasingly greater group size selections (**Fig. 5a**, five middle panels). Strikingly, the 26 reactions with strongly positive Olden Importance scores overlap perfectly with IIMDR values of highest magnitude when centroids from knockouts representing the top and bottom 5^th^ percentile of ΔRLS values are binned. This pattern continues to hold when binning centroids from knockouts out to the top and bottom 12^th^ percentiles of ΔRLS values and is further underscored by the analogous distribution of IIMDR values calculated for the 160 reactions when the 43,000 flux distributions from knockouts belonging to the top and bottom 5th percentile of ΔRLS values are used (**Fig. 5a**, right most panel). Evidently, RNN_536 operates largely by determining whether a centroid flux configuration belongs to one of the same three classes that the CfNN- and CNN ensembles identified (positive, zero (wild-type) and negative ΔRLS). To make its final prediction, RNN_536 must employ other nodes of either less positive or negative Olden Importance to then embellish this initial decision and turn it into a continuous-value function.

### Identity of Reactions Controlling Replicative Lifespan in Yeast

Each of the seven models established in this study identified a set of reactions for use in ΔRLS prediction. The constitution of six of these sets is provided in **Supp. Table K**, along with the composition of every overlap observed between the sets, and the associated hypergeometric probability for randomly observing such an overlap. We have excluded CNN_A to reduce the complexity of the analysis. For PCA_cen and PCA_bin, the statistical analyses we provide in **Supp. Table K** uses the 1305 and 824 reactions that have loading scores residing in the top or bottom 5^th^ percentile of at least one PC driving the respective models. There are 57 possible ways a reaction can be shared among six models. We observed reactions filling all combinations. Presented in **Fig. 5b** (*upper panel)* is a proportionate Euler diagram summarizing the distribution of shared reactions among the six models. A second Euler diagram is presented in **Fig. 5b** (*lower panel)* showing the distribution of shared reactions among the six models when a smaller reaction set for PCA_cen and PCA_bin is employed (comprised of 593 and 555 reactions, respectively). The smaller set of reactions for PCA_cen and for PCA_bin was extracted from the PCs driving each model after all loading scores were weighted using the flux difference between mutants belonging to the top and bottom 5^th^ percentile of ΔRLS values and an empirically determined product-score cutoff was applied (see **Fig. 2n**). It is clear from both Euler diagrams that all six models show remarkable overlap. For 55 of the 57 possible ways reactions could be shared between the six models, the number of reactions shared differed significantly from chance expectation (hypergeometric probabilities ranged from 0.0033 to zero, **Supp. Table K**). Only for overlaps between RNN_536 and PCA_cen, and RNN_536 and CfNN_2009, was the number of reactions observed (N = 54 and 22, respectively) not different from expectation.

One caveat in using hypergeometric probability calculations to assess the overlap between groups that involve greater than two populations, is the order of successive pairwise-testing influences the final probability. We therefore established a second method to determine the significance of the overlaps which we observed between the reactions identified by each model. Briefly, we simulated the starting reaction composition of the six models and then randomly withdrew combinations of reactions equal to the number of reactions required for each model’s accuracy. We did this 100,000 times, and for each of the 57 possible ways to share reactions among the six models we established frequency distributions that quantified the number of reactions observed in each type of overlap. Using these distributions, we assigned a probability to the actual observed number of shared reactions. This form of significance testing addresses a question that is slightly different from the one answered using the hypergeometric probability distribution. Namely, unlike hypergeometric probability testing where the composition of one model’s hits is always assigned, all reaction selections in this sampling approach are randomly chosen at the start of each iteration. Using this arguably more stringent test, we determined that 35 of the 57 possible types of overlap that we observed, still contained many more reactions than expected by chance (**Supplemental Fig. J**).

How can it be that most of the six ΔRLS models share significantly more reactions than expected by chance yet visually, as seen by assessment of the post-IGTD reaction location maps shown in **Figs. 3h, 4f** and **5a**, and from the Euler diagrams of **Fig. 5b**, there are clear distinctions in the reactions that drive each model? The exciting answer to this question is provided in **Fig. 5c**. Up to this point we have assessed the contribution of individual reactions to the predictive power of each of the six ΔRLS models using metrics specific to each modeling technique, making it hard to compare results across the separate studies. So, to do such a comparison, we established a uniform metric. First, for PCA_cen and PCA_bin we took the 593 and 555 product scores of the reduced reaction sets and range scaled them between 0 and 1. For RNN_536 we took the absolute values of all 160 nonzero Olden Importance Scores and similarly range scaled them between 0 and 1. For CfNN_526 and CfNN_2009, all non-zero ‘percent reduction in ensemble accuracy’ scores were converted to positive values then range-scaled between 0 and 1. And lastly, for CNN_B, all ‘percent reduction in ensemble accuracy’ scores > 5% were range-scaled between 0 and 1. This resulted in a total of 1,743 reactions. Focusing only on those reactions that were important to more than one model (908 in total), we assigned the range-scaled score that corresponded to the highest measured value across all relevant models and then plotted this subset of reactions onto the 54×54 reaction location matrix generated by the IGTD algorithm during CNN model construction (**Fig. 5c**, *left panel*). Most of the reactions are evidently of small impact and vanish into the background. Two broad clusters of reactions are, however, evident (*bracketed*). They are more easily discernable using a scatterplot of the range-scaled scores versus a linear ordering of the reactions (54×54 row concatenation) (**Fig. 5c**, *middle panel, asterisks)*. We selected the highest-ranking of these reactions in both clusters (range-scale score > 0.6), then mapped the location of the resulting 90 reactions onto the post-IGTD reaction correlation matrix, which was initially derived using the 43,500 flux configurations from knockouts with ΔRLS values representing the extreme 5^th^ percentiles (**Fig. 5c**, *right panel, arrows)*. Notably, the two clusters of reactions are strongly anti-correlated. Evidently, all six ΔRLS prediction models recognized the same changes in metabolism to predict replicative lifespan – except that either of two sets of reactions were chosen to do so because the sets are anti-correlated with each other.

To map the finer details of the relationships among the highest-ranking 90 reactions, we extracted all the correlation coefficients among these reactions from the post-IGTD correlation matrix shown in **Fig. 5c**, (*right panel),* and then used t-distributed stochastic neighbor embedding (tSNE) to cluster the resulting 90×90 vector profiles (**Fig. 5d**). We then used these groupings to construct a heat map based on correlation coefficient magnitudes **Fig. 5e**. Notably, the two reaction clusters which we identified in **Fig. 5c** remained largely unchanged despite the obvious appearance of finer relationship structure (**Fig. 5e**, *colored text)*. To determine how the flux rate of each reaction varied with replicative lifespan, we again used the 43,500 flux configurations associated with ΔRLS values representing the extreme 5^th^ percentiles. We constructed boxplots for each of the 90 reactions to summarize the flux differentials between knockouts with the highest and lowest ΔRLS values (**Supp Fig. L**). Reactions belonging to each tSNE group displayed similar profiles, so we selected the reaction in each group with the largest range-scaled score differential for presentation in the main figure (**Fig. 5f**, *box plots*). Focusing on the notched region in each box, (notches that do not overlap have different medians at the 5% significance level), we see that flux readings for reactions in tSNE Groups 1 and 5 are significantly higher in yeast knockouts with the highest ΔRLS values versus those with the lowest. A reversal of this flux differential is observed for reactions in tSNE Groups 2, 4, and 6. Among the reactions controlling replicative lifespan, it is noteworthy we did not identify the reaction used to represent biomass production in the Yeast 8.5.0 model (**Fig. 5g**). This implies metabolic flux configurations that are associated with altered replicative lifespan do not simply slow down or speed up growth, rather, they change the routes by which metabolites are moved through the metabolic network (see Metabolic Map, **Supplemental Data**).

Collectively, our data point toward two exciting conclusions regarding lifespan determination in yeast. First, each metabolic flux configuration appears to be associated with an inherent survival probability. In other words, length of life is an emergent property of metabolic flux configuration. Second, although there is an infinite number of flux configurations available to the yeast metabolic network (albeit constrained within physicochemical bounds), there appears to be just three pseudo-metabolic states that these configurations converge toward, and each one is associated with a major shift in survival probability (**Fig. 5h**).

## DISCUSSION

As rousing as our conclusions are regarding life span determination in yeast, we must temper them and acknowledge that the Yeast 8.5.0 genome-scale metabolic model employed in this study, represents just one iteration of a community model undergoing continuing refinement. Indeed the latest GEM, Yeast 9.0, has just been released^14^. Despite its remarkable complexity, the Yeast 8.5.0 GEM contains an incomplete description of lipid metabolism (as does Yeast 9.0). Moreover, the rules that this GEM uses to determine enzyme redundancy rely, for the most part, on protein homology, but recent studies show that incorporating mRNA co-expression data leads to more accurate predictions. Lastly, for some reactions, flux bounds are knowingly broad. Newer tools that measure enzyme concentrations or estimate enzyme *kcat* values continue to refine these ranges.

In this light, the results obtained by our study should be viewed as the first step in an evolving research endeavor. Clearly, the metabolic configurations that we have predicted for each knockout mutant are approximations at best. That said, Yeast 8.5.0 and GEMs like it, continue to be used extensively in both the academic and industrial settings, and their worth in predicting changes to metabolic flux following different kinds of genetic alteration are not disputed. Just as the GEM field has made fantastic strides in iteratively refining the reconstruction of the metabolism of *S. cerevisiae*, we expect that the techniques that we have developed in this paper will serve as a foundation for using metabolism to predict lifespan in yeast and other models for which GEMs exist. As more information is incorporated into GEMs, we anticipate that the relationship between aging and metabolism will be further refined. Whether it is true that lifespan is tri-stable or n-stable, in yeast or any other organisms, remains an exciting possibility. During our study, we note that a GEM for *S. cerevisiae* that incorporate enzyme concentration (ec) constraints was introduced and, in turn, successfully used to alter heme production^15^. Also, as mentioned, Yeast 9.0 was released a couple months ago^14^, and this model contains an additional 73 reactions, 62 metabolites and 13 genes, as well as newly added thermodynamic information for standard biochemical conditions. Notably, deep learning approaches have been used to predict enzyme *kcat* values to further enhance the ec-GEM model^16^. Finally, there is an end game in the GEM field to incorporate both DNA transcriptional control and RNA expression data to have a fully integrated synthetic model of cellular life (multi-scale models)^17^. Techniques that we introduce in this study also have room for improvement. For example, we are yet to devise approaches that scan for optimal assignments of ΔRLS values to individual flux configurations. For computational and cost reasons, the approach we relied upon assigned a single centroid value to all the flux configurations of each knockout.

Notwithstanding the above limitations, can we gauge the relative reliability of our current predictions? In our effort to answer this question, we were shocked to discover that 359 of the 812 single-gene knockouts used in this study (44%) had flux space assignments identical to wild type. It is immediately clear that this observation likely explains many of the modeling challenges that we faced during our investigation. It also raises an additional flag: Our working hypothesis was survival potential is dependent upon metabolic configuration, and so we would have predicted that the replicative lifespan of these mutants would be the same as wild type, but of course we know this is not true. There is nothing extraordinary about the RLS values of the 359 KOs. We find they are scattered randomly across the full range of replicative lifespans that we measured (Supp Table 1). We conclude then, either our working hypothesis is wrong and there are 359 metabolic enzymes that moonlight outside of metabolism in some unknown manner to affect survival, or the flux space assignments made by the Yeast 8.5.0 GEM for these 359 KOs are inaccurate. Almost certainly the latter is true. Shortly, we shall show that although these incorrect flux space assignments did impact the final list of reactions found by our modeling approaches, the lists were augmented with a subset of random reactions that can be identified and excluded, leaving the real signal.

Specifically, the CNN_B ensemble of convolutional neural networks achieved 100% accuracy when distinguishing the 43,500 metabolic configurations belonging to knockout yeast with ΔRLS values in the highest and lowest 5^th^ percentiles. Interrogation of the flux space predictions made by the Yeast 8.5.0 GEM for the 87 KO mutants representing this set of yeast reveal that 44 of the mutants (51%) occupy the same flux space as wild type. How then did the CNN_B ensemble achieve 100% accuracy? Analysis of Supplemental Table K clarifies this answer. It appears that during training, the CNN_B ensemble learned to distinguish four groups of metabolic configurations instead of only two, as we originally believed. That is, the CNN_B network learned to segregate KOs from the two ΔRLS extremes that had real configurational differences by using reactions that truly differed between them. Simultaneously, the CNN_B network learned to distinguish KOs that were assigned wild type configurations in both groups, by using reactions that, through chance alone, had median differences that were exploitable. Recall that we used 500 configurations per KO which were randomly sampled from their FVA-defined solution space. The choice of which reactions CNN_B used to distinguish the second pair of KOs was ultimately decided at this step, specifically by the random assignment of reaction rates to each reaction. This use of four different groups becomes evident when IIMDR values are considered (refer to **Supplemental File**: SUPP_TABLE_medians_and_IIMDRs_cluster1and2_all).

Based on the process of reaction selection used by the CNN_B ensemble, we hypothesize that the reactions contained in Cluster 1 and Cluster 2 of **Fig. 5c** are likely also of two types – real and artifactual. This is because both clusters were identified by combining the results from six separate modeling approaches that all drew from the same set of 406,000 configurations. There are two approaches that theoretically could be used to differentiate both reaction types. First, if a new set of 500 configurations for each KO is generated and our entire set of modeling analyses were to be repeated, we would expect a new subset of artifactual reactions to replace the old ones, since they are ultimately randomly generated. The real reactions, however, should reappear in the new analysis. Second, if our entire analysis were to be repeated using only KOs whose flux solution space is demonstrably different from wild type, only real reactions should reappear in the new analysis. We tested the latter approach, directing our attention to just the 87 knockouts representing yeast with the highest and lowest 5% ΔRLS values. In this set of KOs, there are 43 mutants that the Yeast 8.5.0 GEM predicts have flux spaces that differ from wild type. Since all 87 mutants have a life span phenotype, we confidently updated the redundancy rules for the subset of mutants whose gene products only affected a single reaction but were initially defined as being redundant with one or more other enzymes, but obviously were not. An additional 10 mutants were therefore identified whose flux space differed from wild type when the flux rate through the single reaction they catalyzed was set to zero (or in two cases, when reduced to a level that caused the growth rate to be 10% of wild type). Finally, 10 of the 45 mutants corresponding to yeast with ΔRLS values in the lowest 5^th^ percentile were vacuolar membrane (H^+^)-ATPase (Vma) mutants. We selected only one of these, reducing any bias that might have driven the original analysis results. Using a final collection of 44 mutants then, represented by 22,000 unique metabolic configurations, we constructed new CfNN and CNN models, isolated the reactions responsible for their accuracy, then compared the resulting reaction sets with the reactions comprising Clusters 1 and 2 (Supplemental Data). Significant overlap between the datasets was observed. Of 48 reactions that could have been identified among a set of 2473, 18 reactions in total were re-discovered (sequential hypergeometric probability, *p < 0.004)* (Supp File: non-zero CNN_2473_hits_comparisons). This limited analysis further underscores our initial conclusions, that length of life is an emergent property of metabolic flux configuration, and that there is a convergence among metabolic configurations toward meta-stable flux states. As newer GEMs become available it will be exciting to repeat our analyses in full.

Final mention should be made of the IIMDR metric that we introduced in this study. Initially established as a tool for assessing the quality of our neural net modeling, we note that this metric was also surprisingly effective at predicting which variables would ultimately be employed by our fully trained networks for object discrimination. Use of IIMDR values by the broader artificial intelligence community might therefore be useful for reducing the complexity of training data to speed up the training process.

## Supporting information

Metabolic Map

Main Figure Legends

## ACKNOWLEDGEMENTS

T.A.U.H. was supported by NIH and NIA award F31AG062112, by NIH/NIA training grant T32AG000266-24, and RF1 AG 068908-02. NIH grants R56AG038688 and R21AG054121.

## AUTHOR CONTRIBUTIONS

T.A.U.H. and S.L.R. designed research, performed research, analyzed data, provided Matlab, R, and Python code, and wrote the paper; S.L.R. provided experimental guidance and funding.

## DATA AVAILABILITY

The code used for this project can be found in the github repository yeast-GEM-Lifespan-Modeling (https://github.com/thilsabe/yeast-GEM-Lifespan-Modeling). Data files generated using this code can be found at the figshare repository Hilsabeck, et. al. 2024 - Deep Learning of Cellular Metabolic Flux Distributions Predicts Lifespan (https://figshare.com/s/4e034dea2e6822a8aa6a).

---

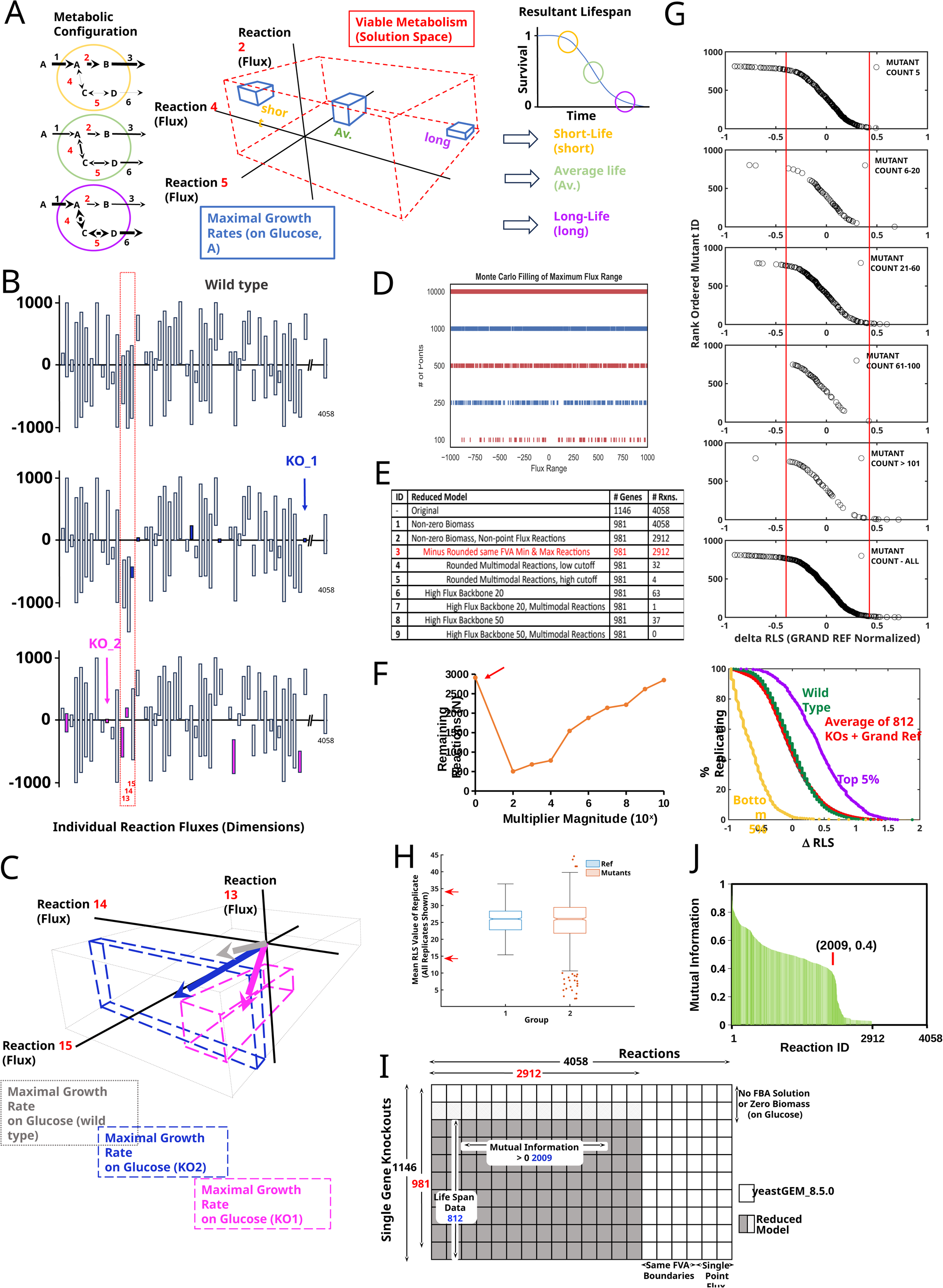

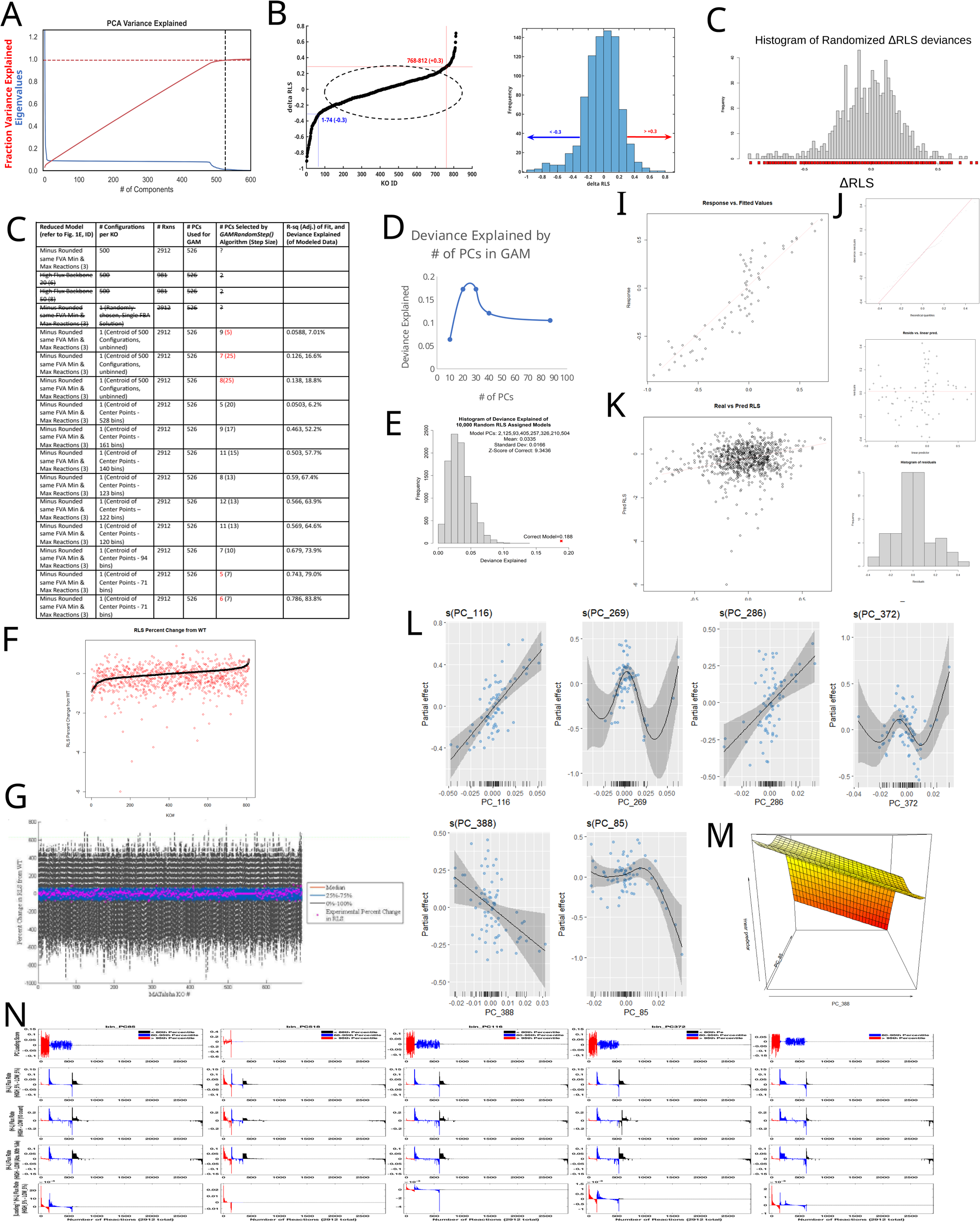

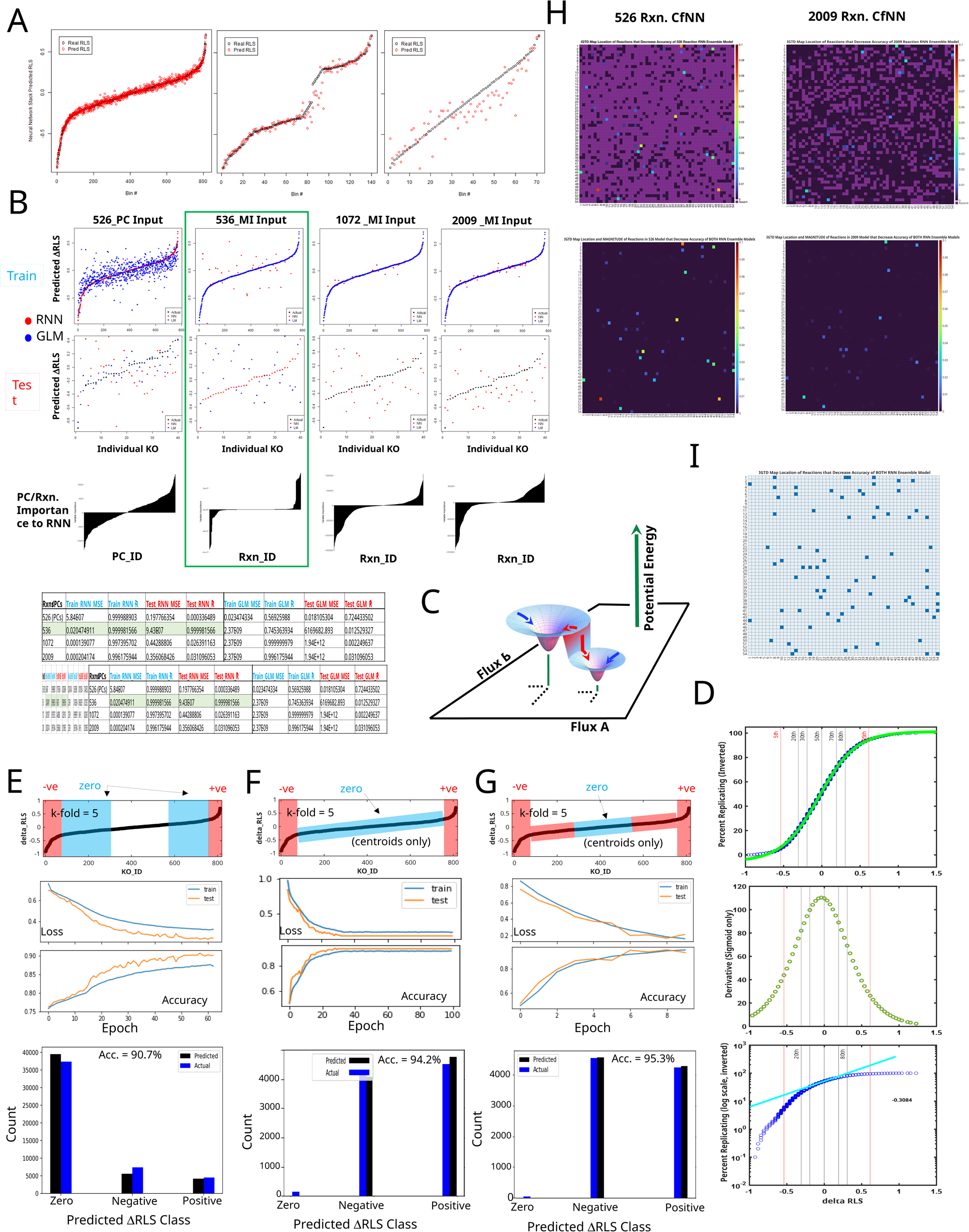

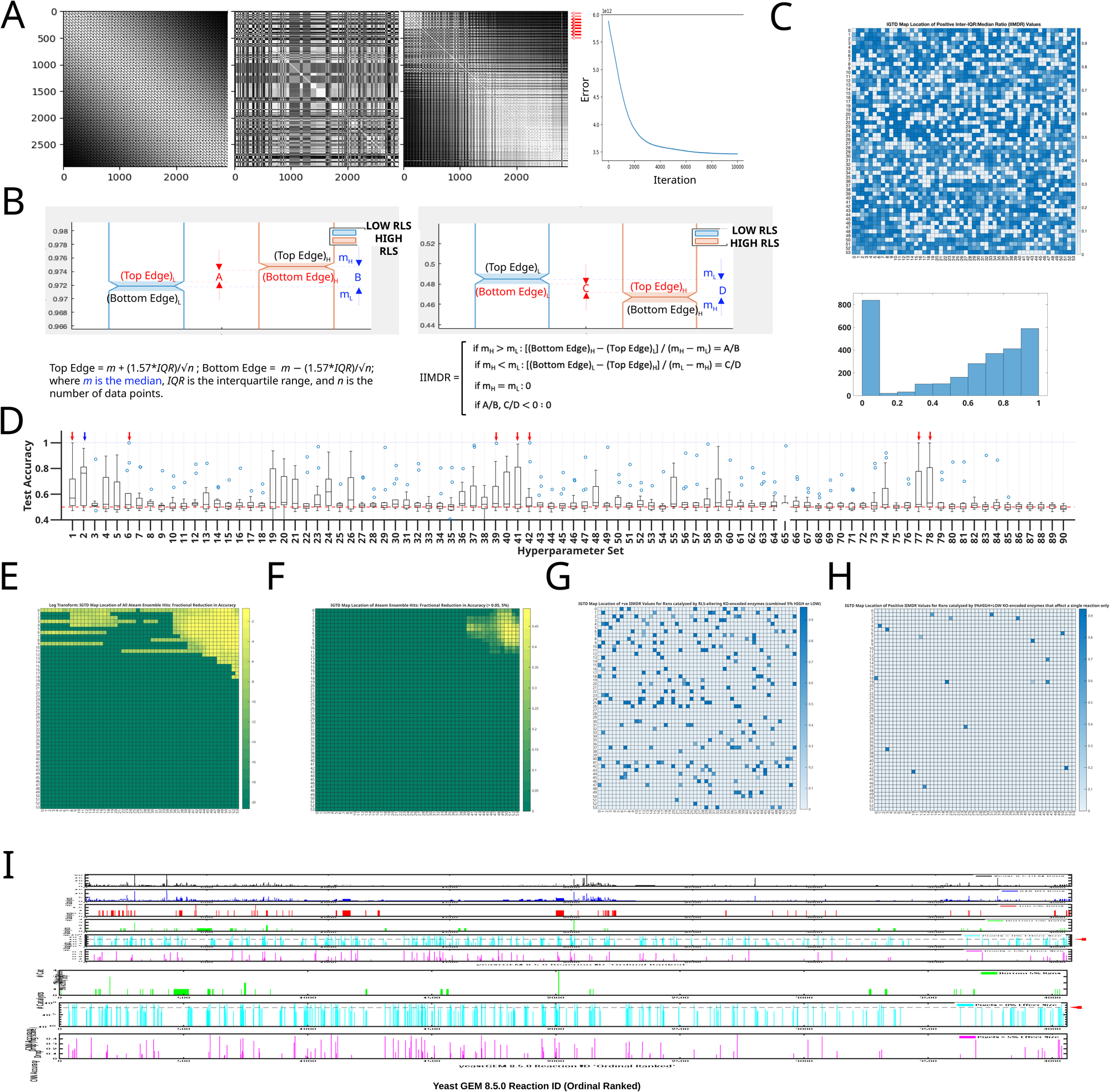

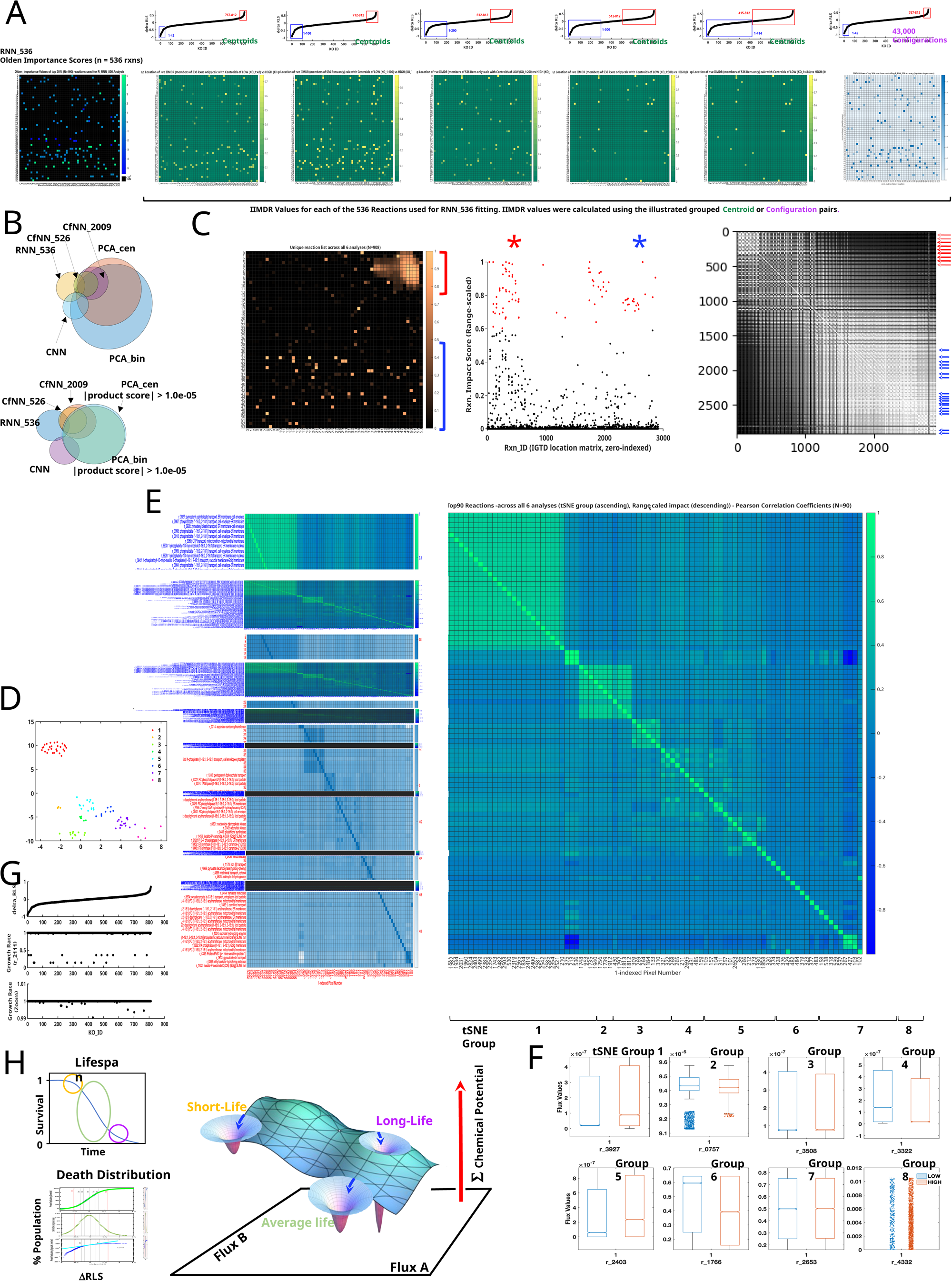

