## Supplementary figures and images for "Deep Learning of Cellular Metabolic Flux Distributions Predicts Lifespan"

### Metabolic Map

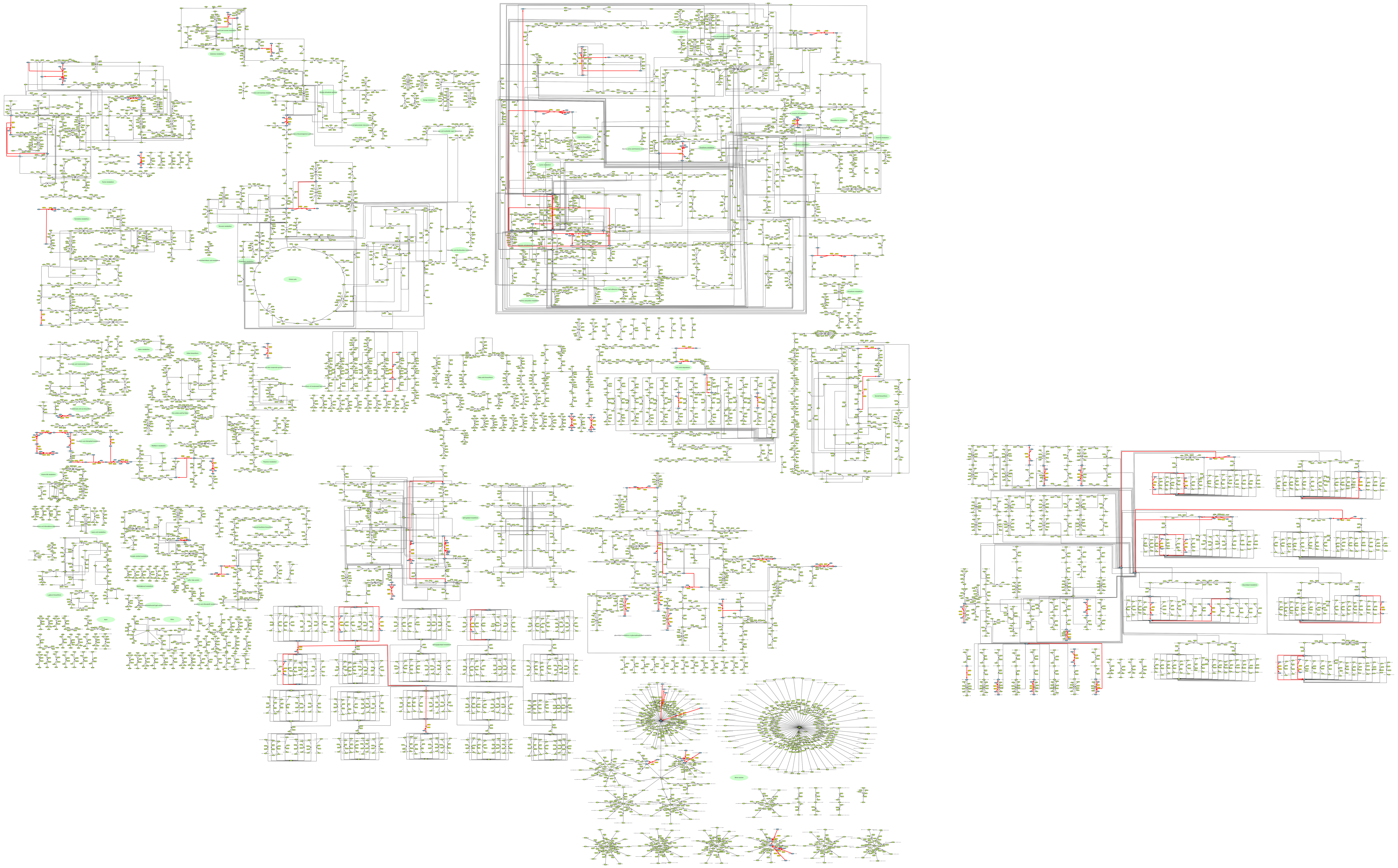

### Supplemental Figure J

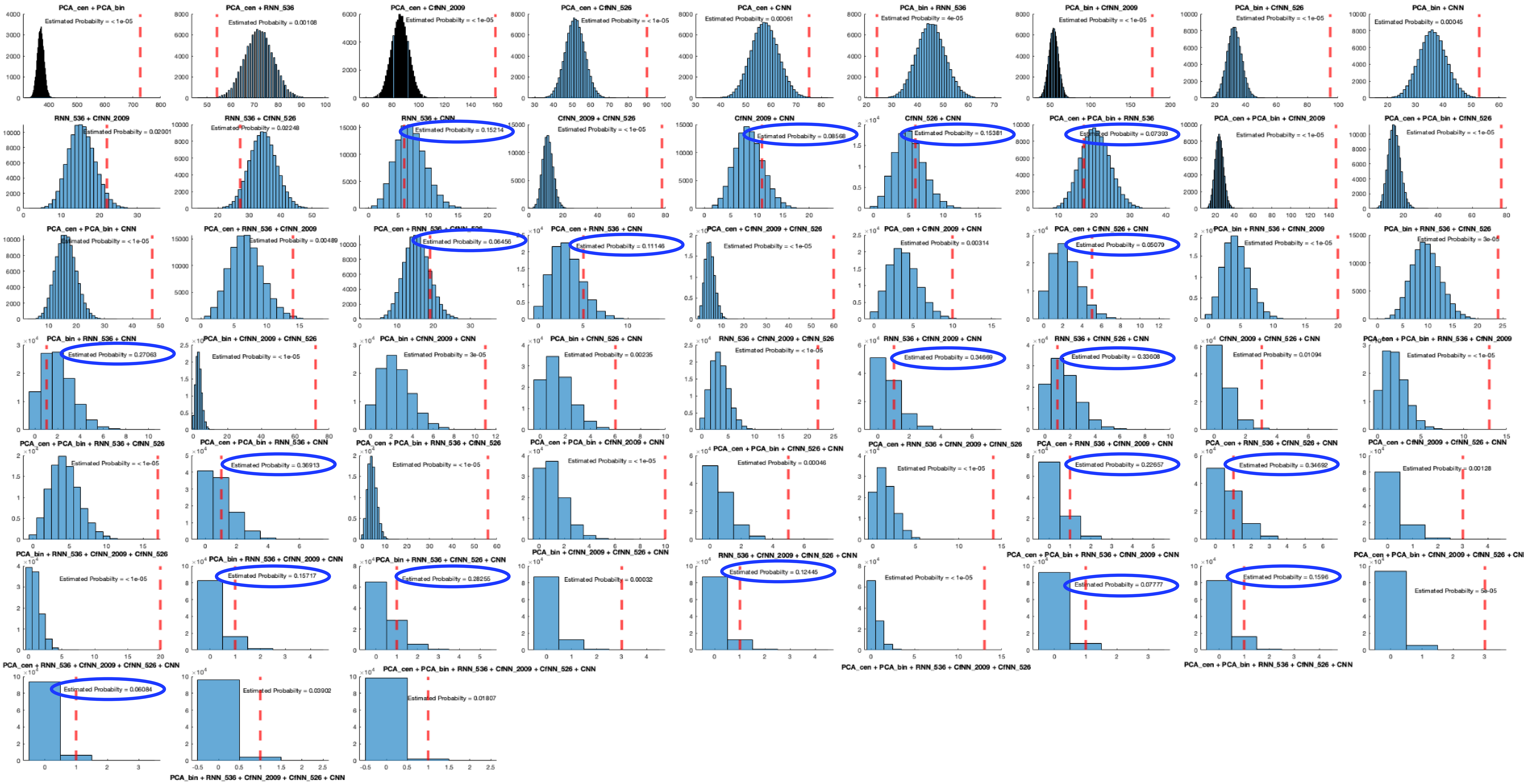

### Supplemental Figure L

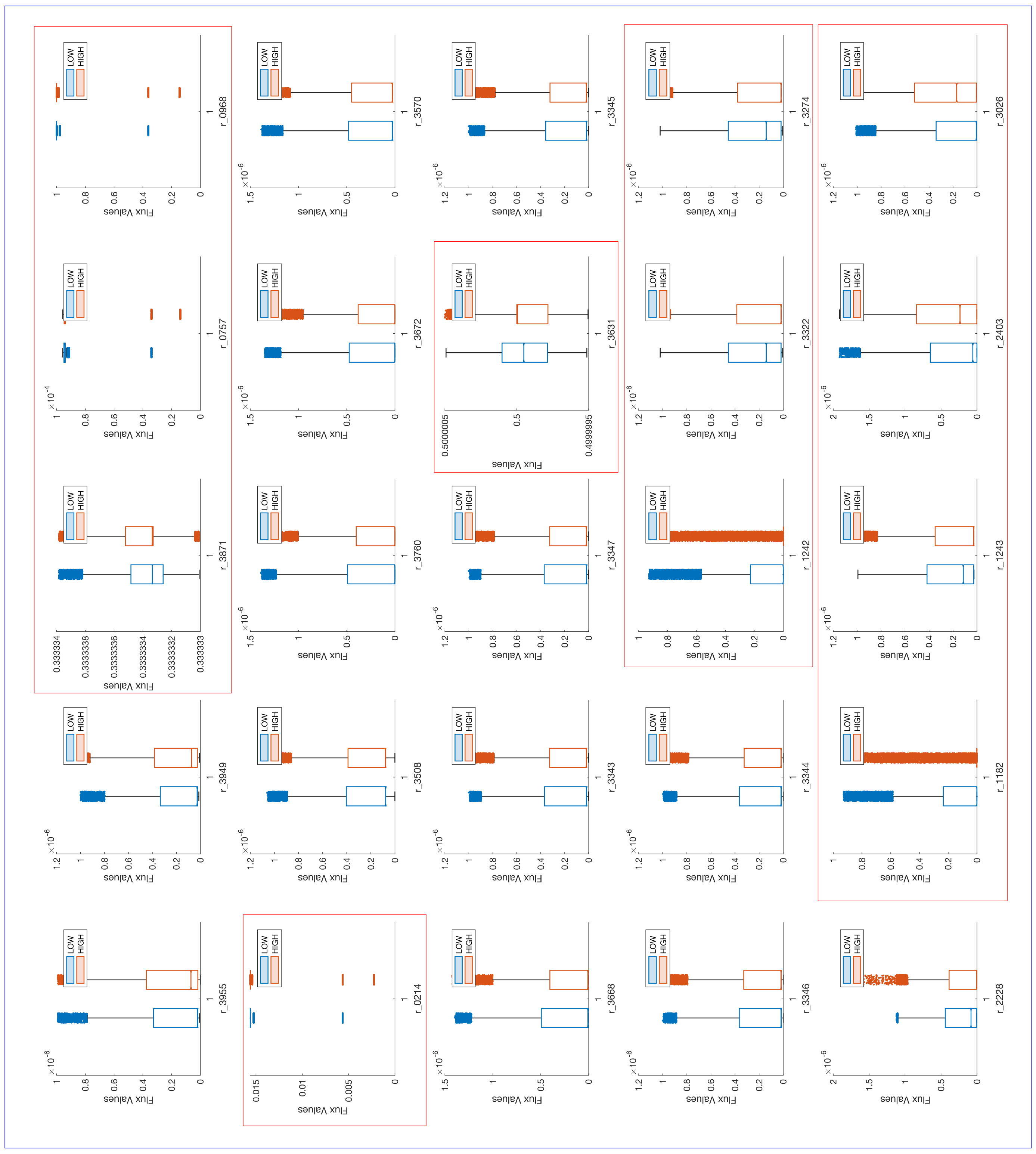

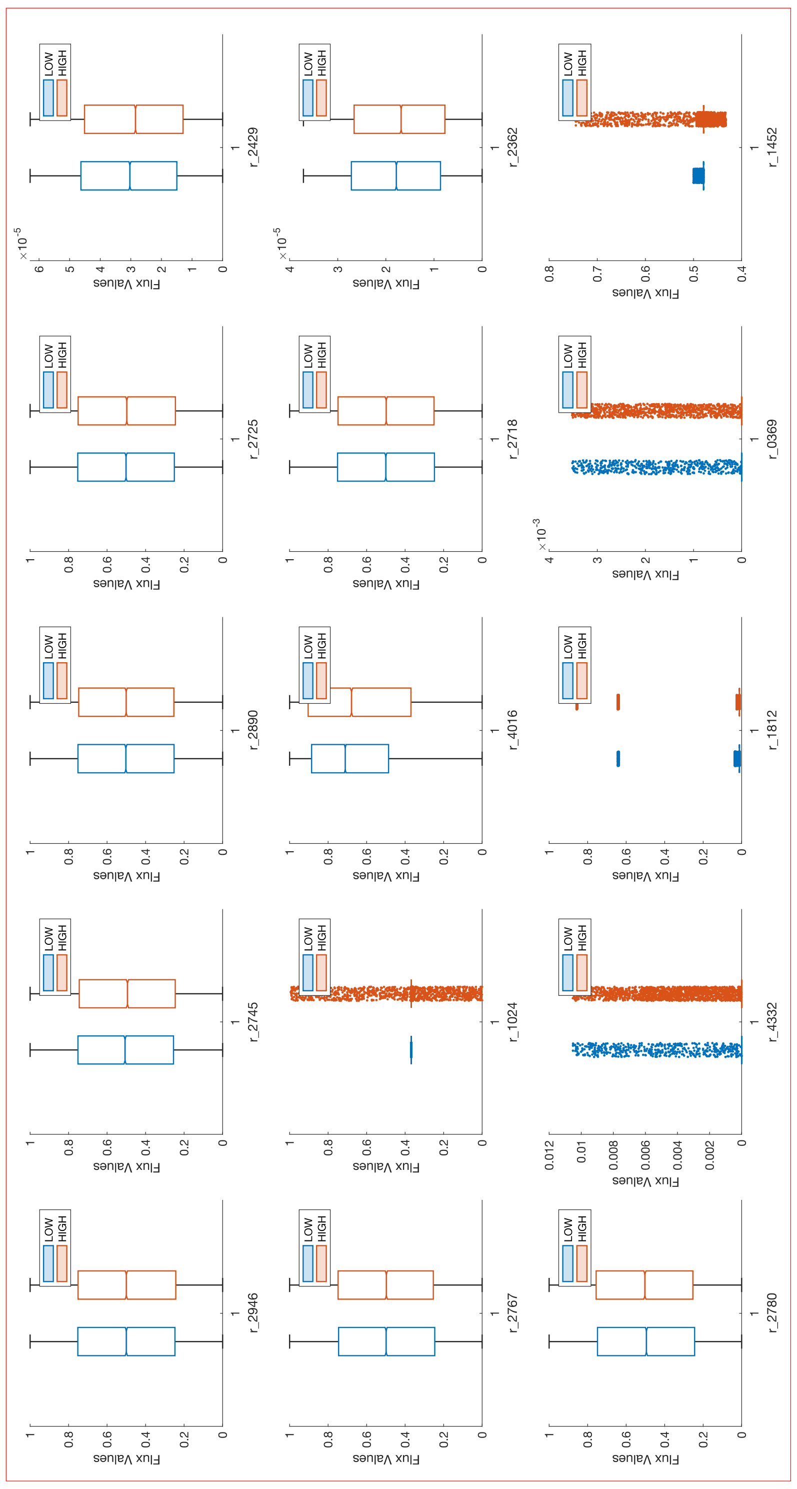

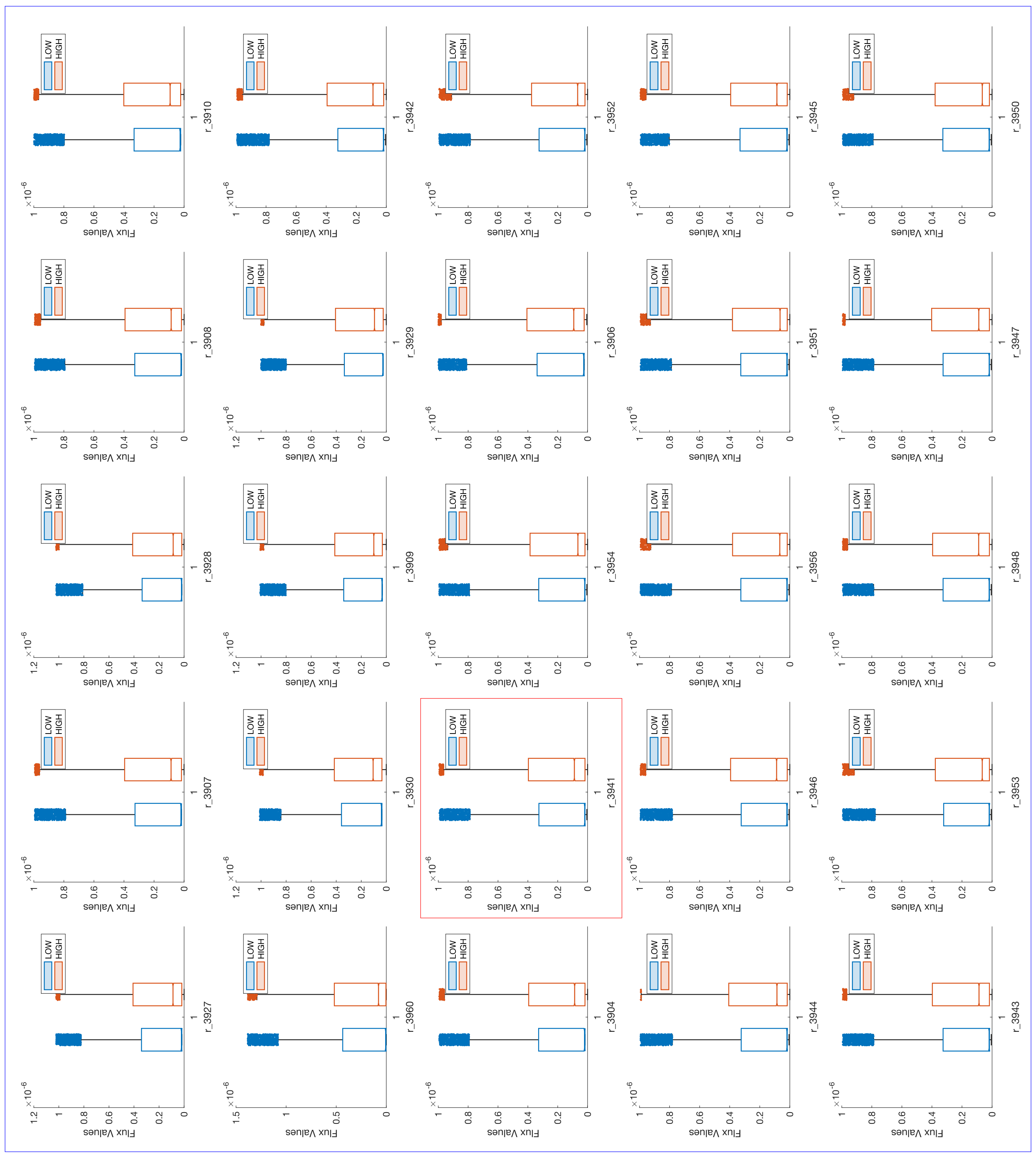

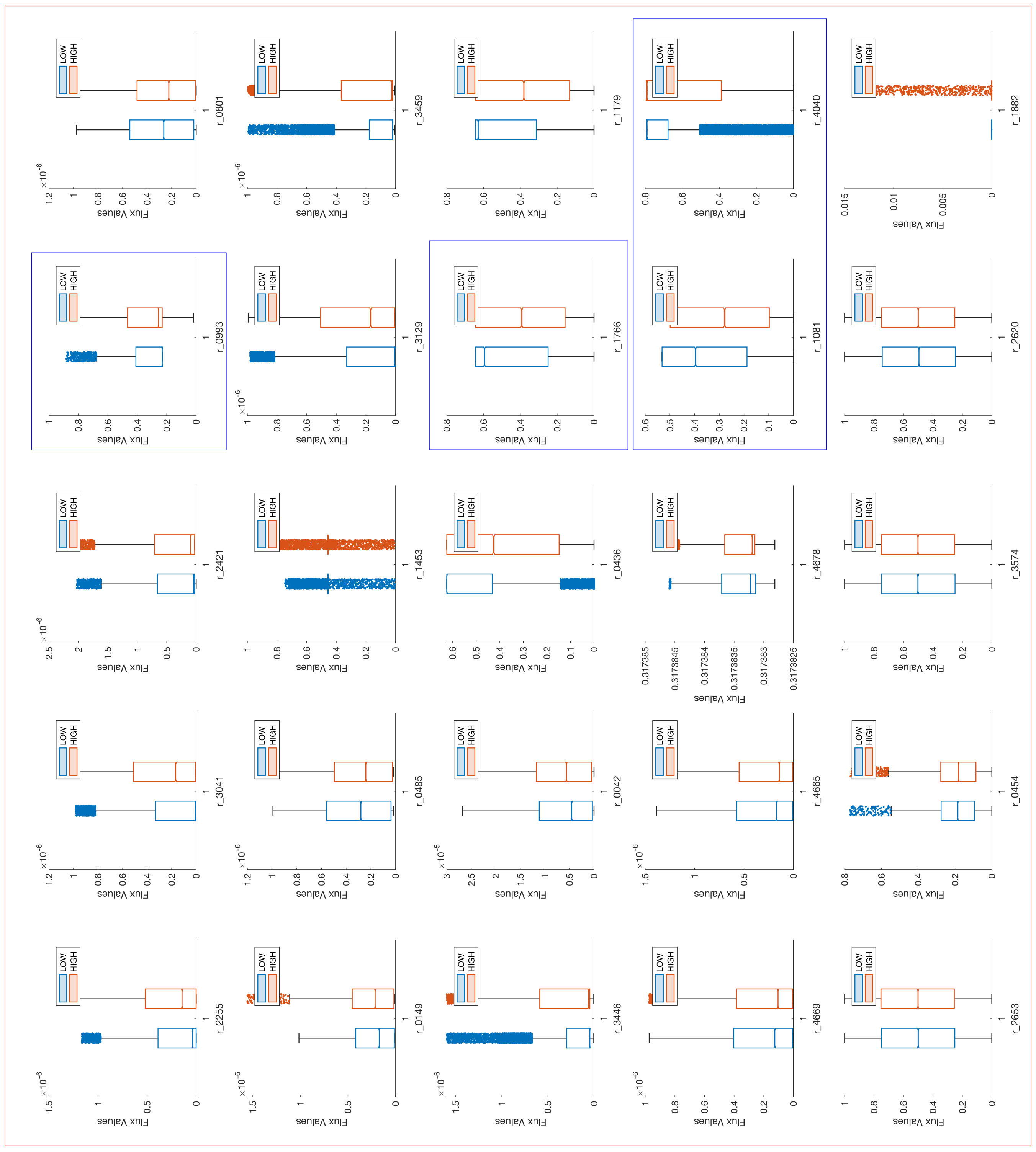
