## Supplementary material for "Deep Learning of Cellular Metabolic Flux Distributions Predicts Lifespan": Main Figure Legends

**Figure 1**

LEGEND:

A. Hypothesis: Redundancies in the reaction network of intermediary metabolism means genetically identical yeast grown under identical growth conditions are not the same. Schematic illustrates hypothetical effect on maximal growth rate and lifespan of tri-stability at a glucose metabolic branch point.

B. Flux variability analysis (FVA) determines the upper and lower limits of every reaction when maximal growth rate on glucose is simulated using the yeastGEM_8.5.0 model of intermediary metabolism.

C. RLS measurements for knockout mutants of each gene contained in yeastGEM_8.5.0, permit exploration of the relationship between RLS and metabolic space. By removing all reactions catalyzed by a KO gene product, the viable reaction flux space is constrained. This, in turn, constrains the flux space associated with maximal growth on glucose (boxes). Arrows point to the center point of each box and highlights that, even though the three boxes overlap, there is an increase in mutual information since the probability of having a discrete type of metabolic configuration is no longer the same as wild type.

D. Monte Carlo sampling was used to determine the optimal number of metabolic configurations required to represent the solution space of each mutant. Five sampling distributions are shown for a decreasing number of generated configurations. Each distribution randomly spans the maximal reaction flux range (+/- 1000 mmol gDW-1 h-1) used in the model.

E. Summary of process used to reduce the number of reactions ultimately used for model construction.

F. Further refinement of reaction reduction process. To remove reactions with near-zero flux values, a multiplier was applied before subtraction of the absolute values of the FVA minima from their corresponding maxima. Arrows mark final chosen cutoff for reduced model choice.

G. Summary of yeast replicative lifespan data used for this study. RLS data is normalized relative to wild type yeast and is presented as delta RLS. For wild type and mutant yeast, since multiple RLS experiments were collected for each, to obtain a final RLS value the mean of each replicate was weighted by group count before pooling and plotting. Pooled data for wild type yeast (green curve), or wild type yeast plus the 812 KO mutants that were predicted to be viable on glucose by yeastGEM_8.5.0 (red curve). Pooled RLS curves for mutants exhibiting the greatest (top 5%, purple curve), or the least (bottom 5%, orange curve) delta RLS values, on average, are also shown.

H. Boxplot of Means from each RLS assay - Mut vs Corresponding Control. Arrows indicate mean RLS values above which mutants that were selected in the 95th and 5th percentiles fall. These correspond to the 98.2th percentile and 0.12% percentiles of wild type data. Whiskers (Q3+1.5*IQR, Q1-1.5*IQR, IQR = Q1-Q3)

(For more information on how the quartiles are computed, see quantile, where the upper quartile corresponds to the 0.75 quantile and the lower quartile corresponds to the 0.25 quantile.

Outliers are values that are more than 1.5 · IQR away from the top or bottom of the box. By default, boxchart displays each outlier using an 'o' symbol. The outlier computation is comparable to that of the isoutlier function with the 'quartiles' method.

The whiskers are lines that extend above and below each box. One whisker connects the upper quartile to the nonoutlier maximum (the maximum data value that is not an outlier), and the other connects the lower quartile to the nonoutlier minimum (the minimum data value that is not an outlier).

Notches help you compare sample medians across multiple box charts. When you specify 'Notch','on', the boxchart function creates a tapered, shaded region around each median. Box charts whose notches do not overlap have different medians at the 5% significance level. The significance level is based on a normal distribution assumption, but the median comparison is reasonably robust for other distributions.

The top and bottom edges of the notch region correspond to m+(1.57⋅IQR)/√n and m−(1.57⋅IQR)/√n, respectively, where m is the median, IQR is the interquartile range, and n is the number of data points, excluding NaN values.

I-J. Summary schematic showing number of KOs (genes) and reactions used for modeling. Values in red indicate the number of yeast predicted to be viable on glucose medium by yeastGEM_8.5.0, and the number of reactions with variance after FVA. PCA and CNN modeling utilized 2912 reactions, while a subset of these reactions (2009) was employed for RNN modeling. Each of the latter reactions had mutual information values >0 for RLS prediction (supplemental file 1) Note we have deltaRLS values for 890 yeast (meaning at least 78 KOs were wrongly predicted to be lethal on glucose when in fact they are not.

**Figure 2**

PCA-GAM model construction: Initial set of parameter optimizations tested for deltaRLS vs. PC model building. The number of included reactions and metabolic configurations was varied. The Center point model was chosen. Final model explained 30% of the delta RLS deviance.

A. Combined scree (blue) and explained variance (red) plots. The first 526 of 2916 PCs accounted for 99% of the variance in the center point vs delta RLS model.

B. Histograms showing model prediction vs. actual delta RLS values (left, and Z-scaled data, right).

C. Model robustness: RLS values were randomized among center point configurations and delta RLS deviance re-measured. Histogram shows distribution of deltaRLS explained deviance values for 10,000 iterations. Red marks shows position of original model.

D. The centerpoint model was used to predict deltaRLS values for all 500 configurations for each of the 812 KO yeast with known delta RLS values. Yeast with little to no change in delta RLS values unduly weight the centerpoint model.

E. (Left) Segmentation of delta RLS values into three ranges for use in separate centerpoint modeling using the 526 PC parameters. (right) Histogram showing frequency of delta RLS values across yeast KOs.

F. (top row) (Tyler what algorithm was used for binning – left most panels – was it fixed width (yellow) or variable (blue)? Also, for the segmentation, did you optimize the bin number separately for each segment, so they had different counts in each group of bins and then fit each segment to PCS separately? Or did you bin then segment and fit? (toy figures from https://demonstrations.wolfram.com/AutomaticallySelectingHistogramBins/). This is key for understanding why got better model fitting. Optimization of model fitting procedure to include delta G. RLS binning, data segmentation for model fitting, and an expanded set of fitting algorithms. Average – Best single GAM model for each of the three segments, average of predictions using each model for each centerpoint prediction ; Weighted Average – average of 10*the deviance explained for each segmented model prediction; GBM – Stacked, Boosted trees (ensemble of x?? weak learners); GAM – stacked, generalized additive model; GLM – Stacked, Linear model; GLMNET – GLM Stacked, Elastic net. Red box marks best fitting model that was used for reaction extraction from significant PCs.

H. Loading plots (range -1 to +1) showing contribution of each reaction to the top three PCs isolated using the 140 bin, segmented GAM model (red box in G). This is not quite true. Tyler used G as an exploratory tool but was unable to easily extract the PCs from the output. We decided that we would say that H used G for exploration, found the optimal number of bins then went back and repeated his random-sample-exclusion script for finding the PCs that fit delta RLS, that he developed for his thesis.

I. Smoothing functions for 140 bin, segmented centerpoint GAM model. Plots show delta RLS (y-axis) vs. PC value (x-axis) and model estimation (solid line; 95% C.I., dotted line). The x-axis is labeled with the PC number out of the 526 total, y-axis with the eigenvector position out of 2912 as well as the estimated degrees of freedom for the smoothed term on the y-axis.

J. Zoom into region of highest predictive accuracy (right) of corresponding plot in panel I.

K. Average reaction flux values for all reactions in the listed PC for the KOs exhibiting the top 5% count delta RLS values (red) versus those exhibiting the bottom 5% count delta RLS values.

Identity of key reactions driving each PC.

**Figure 3**

A, B. Best-fitting, stacked and unstacked RNN models of all knockout centroid configurations. (A) Stacked model predictions for unbinned centroids (left), centroids of 140-binned centroids (middle) and centroids of 71-binned centroids (right) using 526 reactions with top mutual information as input variables, and a stack size of three. (NeuralNet Stack Unbinned allPC nnet size 100 pred_n10_repeats3_tuneL20_2022-07-22; (middle) NeuralNet Stack Unbinned allPC nnet size 100 pred140_n10_repeats3_tuneL20_2022-07-22; (right) NeuralNet Stack Unbinned allPC nnet size 100 pred71_n10_repeats3_tuneL20_2022-07-22) (B). Four unstacked RNN models using, (L-R), 526 PCs, the top 536, 1072 or 2009 reactions (ranked by mutual information score) as input variables. All unstacked RNNs used the same architecture of 3 hidden layers with 350, 233 and 155 fully connected layers. In (B), plots show each model fit to their training data (N= 772 centroids) (top) and testing data (N = 41 centroids, middle). The effect on model accuracy when each reaction is individually removed is shown in the bottom Olden Score plots. For comparative purposes, the top and middle plots also contain centroid predictions using a General Linear Model, (GLM, blue). Relevant fit statistics for the four unstacked and GLM models are tabulated. Green box indicates the best unstacked RNN model. All RNNs were generated using the R ’neuralnet’ package.

C. Model showing energy vs reaction flux space for three configurations. Bottom of wells represent quantum like flux configuration states where neighboring configurations are drawn back toward (blue arrows). Flux values for only two reactions are shown, but the concept is easily translated to n-space. Moving between flux configurations requires energy input (green arrow).

D. (top) Linear plot of grand Ref lifespan data for all 34,000 wild type yeast. Blue line shows inverted replicative lifespan curve. (middle) derivative of sigmoidal curve fit in top panel showing bell shape of underlying yeast count distribution. (bottom) Log-linear plot of top panel, blue line shows departure from linearity starts at ~20 and 80th percentiles (corresponding to delta RLS values of +/-0.3. We interpret this as lifespan starts to deviate from ”noisy death” and possibly reflects inflection points where there are changes to the average metabolic configuration (ie away from the average flux pathway and its noisy boundaries, perhaps where evolution wants to hold S. cerevisiae) such that the configuration changes have substantive effects on lifespan.

E-G. Three best CfNN models. (Top) Graphic showing source of configurations used in each CfNN model. 500 configurations (fully shaded), the centroid of the 500 configurations (small shading), or no configuration at all was employed for each knockout. The centroid of each knockout is plotted (black line). (Middle), Loss and Accuracy plots during model training for best of five folds. (Bottom), Final test accuracy for best of five folds showing actual vs predicted calls on each configuration.

H, I. (Top row) Heatmaps showing the effect size (ABSOLUTE VALUE) on model accuracy following removal of each reaction in the 525 (left) and 2009 (right) reaction CfNN models. Purple pixels are reactions from the 2912 reduced reaction subset that were not employed in the 526 or 2009 modeling process. Data is present in post-IGTD format, where 2912 (+4 dummy reactions) from the reduced reaction set are arranged into a 2916 pixelated image format that maximizes correlation between neighboring pixels/reactions(see main text). The scale is the same for both panels. No pixel effect size was > 8%. (Bottom panels) Same as respective top panel, except only reactions that negatively impacted BOTH the 525 (left) and 2009 (right) reaction CfNN models when removed are shown. 78 reactions are common. To facilitate recognition of the relevant reactions/pixels, the 78 reactions are highlighted with equal intensity in panel I. Red circle marks location of reactions identified by later described convolutional neural network.

**Figure 4**

An ensemble of convolutional neural networks (CNNs) identifies a core set of coordinated reactions that distinguish long- from short-lived yeast mutants.

A. The IGTD algorithm was used to generate a 54x54 pixel representation of the reduced reaction set (n = 2912), with pixel positions corresponding to the degree of relatedness between reactions in knockout mutants belonging to the top and bottom 5% delta RLS values. Relatedness is defined as the degree of correlation between reaction fluxes across the 43,500 metabolic configurations that span the reaction space of these mutants. The result of the IGTD clustering algorithm is shown. Left panel, Euclidian distance matrix of the pixels in a 54x54 pixel square (this is the target matrix); 2916 feature distance matrix before (middle) and after (right) IGTD algorithm (four dummy reactions were added to the 2912 features before the IGTD algorithm was run, to square the feature distance matrix); (far right) record of the optimization process (horizontal axis is number of iterations, vertical axis is error value). In the matrices, white pixels are more closely related. After optimization, as part of the IGTD routine, each metabolic configuration was mapped onto the reaction location map and pixel intensities assigned. Although 256 grey scale images are presented, image data was stored as double precision, floating point numbers and these values were used for CNN analyses. Arrows mark approximate positions of reactions (rows) corresponding to yellow pixels in panel E (red), and red pixels in Fig. 5A (blue).

B. Definition of the Inter-IQR-to-Median Difference Ratio (IIMDR).

C. IIMDR values for all 2916 reactions, ordered in accord with their IGTD pixel assignment. A histogram quantifying frequency of IIMDR values is shown below.

D. Box plot showing test accuracies for all 90 groups of CNN models. Each group differs by hyperparameter choice (neuron dropout rate (0, 0.1, 0.2), kernel size (3, 6, 9), number of filters (24, 32, 64, 128, and 256) and batch size (16, 32)), as well, 14 different seed choices were used across the groups for initialization of the random number generator. Blue arrow marks the best performing hyperparameter set. The top five models were used from this group to form one ensemble model. The red arrows mark the groups from which the seven top performing models were used to form a second ensemble model.

E. CNN modeling was used to classify metabolic configuration images into two groups – high or low delta RLS. After model building and tuning, an ensemble of the best five predictors was established and testing showed it afforded perfect discriminatory power. Reactions (pixels) driving the success of the ensemble were identified using a dropout approach. Shown are all reactions causing a fractional reduction in accuracy greater than zero. Data are log transformed to aid visualization of the smallest effect sizes.

F. Same as in E, except only reactions that fractionally reduce model accuracy by > 5%, when removed are shown.

G. IGTD map location and IIMDR value (scale) of all reactions in the reduced reaction subset that are catalyzed by an enzyme product which is removed by one of the knockout genes present in either the top or bottom 5% of the 812 KO’s. Note, by definition IIMDR is positive or zero, so only reactions with positive IIMDR values are plotted. Note coloured according to IIMDR values and note most are dark yellow – meaning IIMDR value is close to 1 (max). See photochop file for overlap with pixels in panel C. Point is that CNN is not just finding the reactiosn with the biggest IIMDR values.

H. Same as in F, except the marked reactions correspond to genes that catalyze no other reaction other than the one shown. These are not reactions encoded by single genes, rather they are reactions that have>=1 catalysts and each catalyst only catalyzes this one reaction. (note the first figure I had up for F and G were in fact across all 812 KOs – In the 812 set, 398 KOs have positive delta RLS values and 414 have negative delta RLS values with respect to the wild type set to zero. The color in the plot indicates positive IIMDR values. Note also, the number of KOs among the 812 set that encode an enzyme product that UNIQUELY targets one reaction, and no one other of the 812 KO genes target that reaction ,is 257. Not all have positive IIMDR values. Note also the yeastGEM_8.5.0 contains 1150 genes and I only analyzed the 812 subset since the proteins encoded by these KOs encoded all the necessary enzymes needed by the non-spontaneous reactions of the 2916 reduced reaction set.

I. Effect of removing each reaction on CNN ensemble accuracy. Only those reactions that fractionally reduce model accuracy by > 5% are included in the 2nd plot. It is clear from presenting the data in this form that fractional accuracy values < 5% are mostly noise. NOTE THAT THE SCALE VARIES in each plot. Arrow and dotted line mark 1% line in log plot.

**Figure 5**

A. Reactions driving RNN_536 model accuracy differ from those driving CNN model accuracy. Left panel. Heatmap showing Olden Importance scores for reactions used to build the RNN-536 model. 160 reactions (30%) have scores |>0|. Reactions are mapped onto the same 54x54 reaction location matrix generated by the IGTD algorithm for the 43,000 configurations of yeast with delta RLS values in the top and bottom 5th percentiles generated during CNN model construction. (Recall, the position of reactions in this matrix resulted from the IGTD algorithm optimizing the similarity between the correlation matrix of the 2912 (+4) reactions of the reduced reaction and the distance matrix of a 54x54 pixel matrix). Remaining panels, bottom row. IIMDR values calculated using either centroid values or all 500 configurations (right most panel) for the listed knockout groups (top row). Reactions with the highest IIMDR values calculated using the centroid configurations from 24.6% of the yeast knockouts (bright yellow pixels), representing the 100 smallest and 100 greatest delta RLS values, show obvious overlap with the most important reactions driving accuracy of the RNN_536 model (Olden scores in bright green pixels, left-most panel). These data suggest the RNN_536 model, like the CfNN models, identified a set of reactions that distinguishes three kinds of centroids. Additional reactions presumably work to fine tune delta RLS predictions. Intriguingly, only 5 of the 160 reactions with non-zero Olden scores are shared with the top 128 CNN hits (Supp Table X). This number increases to 20 if we include all 365 non-zero CNN hits.

B. Euler Diagram showing the proportionate overlap among each set of important reactions identified by the 6 models described in this study (top panel). Sets PCA_cen, PCA_bin, RNN_536, CfNN_526, CfNN_2009 and CNN contain 1305, 824, 160, 192, 115 and 128 reactions, respectively). In the bottom panel, we have restricted the size of the important reaction sets for the PCA/GAM studies to reactions with absolute product scores > 1.0e-05 (refer to Fig 2n, bottom row), where product score is calculated as AbsPCloading score*(H-L)5%delta_RLS. Under these restrictions, PCA_cen and PCA_bin reduce to 593 and 555 reactions. See also Supp Fig X for overlap counts.

C. (Left panel) Heatmap showing all reactions necessary for the accuracy of one or more of the six models established in this study. For the CfNN_2009, CfNN_526 and CNN ensembles, we analyzed only the top 192, 115, and 128 reactions affecting accuracy. For RNN_536 we used the top 160 reactions ranked by absolute Olden Importance. For PCA_cen and PCA_bin we restricted analysis to 593 and 555 reactions based on absolute product scores. Each scoring metric was range scaled from 0-1. Reactions important to more than one model were assigned a range-scaled score corresponding to the highest measured value, then all reactions were plotted onto the 54x54 reaction location matrix generated by the IGTD algorithm during CNN model construction. Two clusters of reactions are evident (bracketed), which are even better discernable on a scatterplot of the range-scaled scores versus a linear ordering of the reactions (asterisks, middle panel). We selected the top-most 90 reactions (red dots) then mapped the location of both clusters back onto the post-IGTD, reaction correlation matrix derived using the 43,500 flux configurations from knockouts with extreme delta RLS values (+/- 5th percentiles). Red and blue arrows correspond to the position of reactions from the first and second clusters, respectively.

D-F. To assist in understanding and visualizing the relationship between the top-most 90 reactions that control accuracy among the six delta RLS prediction models, we extracted the correlation coefficients between each of these reaction across the 43,000 flux configurations from knockouts with extreme delta RLS values (+/- 5th percentiles). We used tSNE to cluster the reactions based on similarity between their correlation profiles (D), then the resulting groupings were in turn used to construct a heatmap to visualize the relationship between the reactions based on their correlation coefficient profiles (E). In E, reactions in red correspond to reactions of cluster 1 in panel C. tSNE groupings are shown at the base of the heatmap. Further down, boxplots showing representative flux distributions for the reactions in each of the tSNE groupings are provided. (F) Boxplots were calculated using the flux configurations of knockouts from the high (42x500) and low (45x500) delta RLS groups. Line inside box is median (m); top and bottom edges are upper and lower quartiles, respectively, and the distance between them is the interquartile range (IQR). Notched regions correspond to m+(1.57⋅IQR)/√n and m−(1.57⋅IQR)/√n, respectively, where n is the number of data points. Outlier (circles), defined as more than 1.5*IQR from top or bottom of box. Whiskers connect boxes to max/min data that is not an outlier. Boxes whose notches do not overlap have different medians at the 5% significance level.

G. r_2111 is a pseudoreaction that represents growth in the Yeast_8.5.0 GEM. We find no correlation between r_211 centroid flux rate and delta RLS. Lower panel is a magnification of the middle panel.

H. Model depicting the relationship between delta RLS (lifespan), instantaneous chemical potential (integrated across all metabolites in the Yeast GEM 8.5.0 model), and metabolic configuration at steady state in yeast. (Left) Sketch of a wild type S. cerevisiae survival curve and the corresponding distribution of yeast deaths. (Right) Our data suggest three, meta-stable flux configurations exist in wild type yeast populations, and each is associated with an optimal configuration of minimum chemical potential (wells). All metabolic configurations resonate around one of these energy wells. Movement across the boundaries of each meta-stable flux configuration is possible but requires large flux perturbations (energy input). The energy well associated with long-lived yeast is deepest and equates with flux configurations that inherently have fewer types of metabolites with chemical potentials that puts the cell at risk of undergoing spurious detrimental reactions outside of the essential metabolic network.
